# The Inositol Trisphosphate Receptor (IP_3_R) is Dispensable for Rotavirus-induced Ca^2+^ Signaling and Replication but Critical for Paracrine Ca^2+^ Signals that Prime Uninfected Cells for Rapid Virus Spread

**DOI:** 10.1101/2023.08.09.552719

**Authors:** Jacob L. Perry, Francesca J. Scribano, John T. Gebert, Kristen A. Engevik, Jenna M. Ellis, Joseph M. Hyser

## Abstract

Rotavirus is a leading cause of viral gastroenteritis. A hallmark of rotavirus infection is an increase in cytosolic Ca^2+^ caused by the nonstructural protein 4 (NSP4). NSP4 is a viral ion channel that releases Ca^2+^ from the endoplasmic reticulum (ER) and the increase in Ca^2+^ signaling is critical for rotavirus replication. In addition to NSP4 itself, host inositol 1,4,5- trisphosphate receptor (IP_3_R) ER Ca^2+^ channels may contribute to rotavirus-induced Ca^2+^ signaling and by extension, virus replication. Thus, we set out to determine the role of IP_3_R Ca^2+^ signaling during rotavirus infection using IP_3_R-knockout MA104-GCaMP6s cells (MA104- GCaMP6s-IP_3_R-KO), generated by CRISPR/Cas9 genome editing. Live Ca^2+^ imaging showed that IP_3_R-KO did not reduce Ca^2+^ signaling in infected cells but eliminated rotavirus-induced intercellular Ca^2+^ waves (ICWs) and therefore the increased Ca^2+^ signaling in surrounding, uninfected cells. Further, MA104-GCaMP6s-IP_3_R-TKO cells showed similar rotavirus susceptibility, single-cycle replication, and viral protein expression as parental MA104- GCaMP6s cells. However, MA104-GCaMP6s-IP_3_R-TKO cells exhibited significantly smaller rotavirus plaques, decreased multi-round replication kinetics, and delayed virus spread, suggesting that rotavirus-induced ICW Ca^2+^ signaling stimulates virus replication and spread. Inhibition of ICWs by blocking the P2Y1 receptor also resulted in decreased rotavirus plaque size. Conversely, exogenous expression of P2Y1 in LLC-MK2-GCaMP6s cells, which natively lack P2Y1 and rotavirus ICWs, rescued the generation of rotavirus-induced ICWs and enabled plaque formation. In conclusion, this study shows that NSP4 Ca^2+^ signals fully support rotavirus replication in individual cells; however, IP_3_R is critical for rotavirus-induced ICWs and virus spread by priming Ca^2+^-dependent pathways in surrounding cells.

**Importance:** Many viruses exploit host Ca^2+^ signaling to facilitate their replication; however, little is known about how distinct types of Ca^2+^ signals contribute to the overall dysregulation of Ca^2+^ signaling or promote virus replication. Using cells lacking IP_3_R, a host ER Ca^2+^ channel, we could differentiate between intracellular Ca^2+^ signals within virus-infected cells and intercellular Ca^2+^ waves (ICWs), which increase Ca^2+^ signaling in neighboring, uninfected cells. In infected cells, IP_3_R was dispensable for rotavirus-induced Ca^2+^ signaling and replication, suggesting the rotavirus NSP4 viroporin supplies these signals. However, IP_3_R-mediated ICWs increase rotavirus replication kinetics and spread, indicating that the Ca^2+^ signals from the ICWs may prime nearby uninfected cells to better support virus replication upon eventual infection. This “pre-emptive priming” of uninfected cells by exploiting host intercellular pathways in the vicinity of virus-infected cells represents a novel mechanism for viral reprogramming of the host to gain a replication advantage.

## Introduction

Calcium (Ca^2+^) signaling is a cornerstone of cellular communication and is critical for maintaining homeostasis and responding to damage or infection(1). As such, it is unsurprising that many viruses have evolved strategies to exploit these pathways to facilitate virus replication and spread (2). However, the highly interconnected nature of Ca^2+^ signaling has made it challenging to understand how viral and host proteins interact to orchestrate the patterns of Ca^2+^ dysregulation that are observed during infection. One key strategy for virus-induced calcium dysregulation is the expression of viral ion channels, or viroporins, which can conduct Ca^2+^ across host cell membranes (3). The rotavirus nonstructural protein 4 (NSP4) is among the most well-characterized Ca^2+^ conducting viroporins and rotavirus has become a leading model system to study how viruses exploit Ca^2+^ signaling to promote their replication (4). Yet, the interplay between rotavirus-and host-induced Ca^2+^ signaling pathways remain incompletely characterized.

Numerous studies show that NSP4 elevates cytosolic Ca^2+^ levels during infection (5-7). NSP4 is a multifunctional endoplasmic reticulum (ER), transmembrane glycoprotein (8, 9) that increases the Ca^2+^ permeability of the ER by forming a Ca^2+^ permeable viral ion channel (viroporin) (10, 11). The NSP4-mediated release of ER Ca^2+^ in turn activates a host process known as store-operated calcium entry (SOCE) to further elevate cytosolic Ca^2+^ levels (12). This is critical for multiple steps in rotavirus replication, including the activation of the autophagy pathway and assembly of the rotavirus outer capsid protein, VP7 (12, 13). Yet, NSP4 may play both a direct and an indirect role in ER Ca^2+^ release, through interaction with host channels. The two main eukaryotic ER Ca^2+^ release channels are the ryanodine receptor (RyR) and the inositol-1,4,5-trisphosphate receptor (IP_3_R), and while RyR expression is generally restricted to excitable cells, IP_3_R is widely expressed in most cell types (14). Mammals have three IP_3_R genes, each with multiple splice variants that provides nuance to the regulation of IP_3_R Ca^2+^ release and therefore the ability to shape Ca^2+^ signals (14, 15). The IP_3_R channel is activated by IP3 and channel opening is regulated by cytosolic Ca^2+^ in a biphasic manner, such that low cytosolic Ca^2+^ levels potentiates ER Ca^2+^ release and increasing cytosolic Ca^2+^ levels inhibit ER Ca^2+^ release (16). Thus, even though the NSP4 viroporin directly alters ER and cytosolic Ca^2+^, NSP4 Ca^2+^ release may also potentiate IP_3_R activity, increasing the Ca^2+^ dysregulation of host cells.

In addition to NSP4 and IP_3_R crosstalk within infected cells, we recently discovered that rotavirus infection triggers intercellular Ca^2+^ waves (ICWs) through the release of ADP from infected cells and activation of P2Y1 purinergic receptors on neighboring cells. Activation of P2Y1 receptors results in an IP_3_R-mediated ER Ca^2+^ signal and since many ICWs are produced during infection, this significantly increases Ca^2+^ signaling in neighboring, uninfected cells as well (17). Thus, the ER Ca^2+^ store is a critical source of Ca^2+^ signals in both rotavirus infected and the nearby uninfected cells and IP_3_R has the potential to significantly impact the overall landscape of rotavirus induced Ca^2+^ signaling dysregulation.

Thus, the goal of this study is to examine the contribution of IP_3_R to rotavirus Ca^2+^ dysregulation and by extension, its role in rotavirus infection, replication and spread. While HEK293 and HeLa cells with genetic knockout of all three IP_3_R genes (e.g., IP_3_R1, IP_3_R2, and IP_3_R3) have been established previously, these cell lines are suboptimal for studying rotavirus replication and it is not known whether they support the rotavirus ICW signaling pathway (15, 18). Thus, we generated an IP_3_R triple knockout in MA104 cells, a vervet monkey (*Chlorocebus pygerythrus*) kidney cell line commonly used to study rotavirus (17, 19, 20). We found that IP_3_R was not necessary for aberrant Ca^2+^ signaling in infected cells, but was necessary for Ca^2+^ dysregulation in neighboring, uninfected cells. In parallel, IP_3_R was dispensable for single-round rotavirus infection and replication, but lack of IP_3_R, and therefore lack of ICW signaling, strongly reduced rotavirus spread. This study provides new insights into mechanisms exploited by viruses to reprogram infected cells but also to pre-program surrounding cells for subsequent rounds of infection.

## Materials and Methods

### Cells and viruses

All cells were cultured in complete DMEM [high glucose DMEM supplemented with 10% fetal bovine serum (FBS) and Antibiotic/Antimycotic (Invitrogen)] and incubated at 37°C in 5% CO_2_. HEK293 and HEK293-IP_3_R-TKO cells were provided by Dr. David Yule (15, 21). The MA104 (vervet monkey kidney), LLC-MK2 (rhesus monkey kidney), HEK293 and HEK293-IP_3_R-TKO cell lines were engineered to express the cytosolic Ca^2+^ sensor, GCaMP6s, as previously described (20, 22). All rotavirus infections were at the indicated MOI for 1 hr with a recombinant SA11 expressing mRuby3 from gene 7 (SA11-mRuby), which was propagated as previously described (20, 23).

### MA104-GCaMP6s-IP_3_R triple-knockout generation

#### CRISPR/Cas9 development and transduction

MA104 cells lacking IP_3_R expression (MA104-GCaMP6s-IP_3_R-TKO) were generated by lentivirus transduction to introduce Cas9 and small guide RNAs (gRNAs) to IP_3_R1, IP_3_R2, and IP_3_R3 (Table 1). The lentivirus construct was designed by our lab and manufactured by VectorBuilder (Chicago, IL). MA104-GCaMP6s cells were transduced with MOI 10 in complete DMEM supplemented with 10 µg/mL polybrene (21). At 72 hours post-transduction, cells were passaged in the presence of 40 µg/mL blasticidin. After 2 weeks of selection, cells were dilution cloned and resulting clones were screened for lack of agonist-induced IP_3_R Ca^2+^ responses, as described below.

**Table 1.**
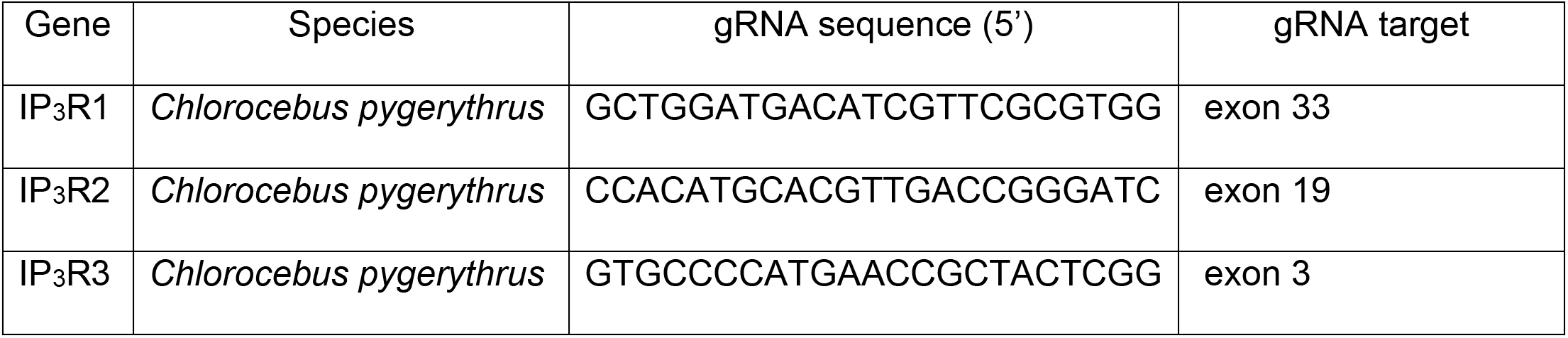
African green monkey IP_3_R1, IP_3_R2, and IP_3_R3 gRNA sequences and exon target sites.

#### *Sequencing of* IP_3_R *triple-knockout*

Genomic DNA was extracted using a PureLink^TM^ gDNA mini kit (Invitrogen, USA). PCR amplification was performed using KOD Hotstart polymerase kit (EMD Millipore) and primers flanking the gRNA target sites (Table 2). PCR products were cloned in TOPO-TA vectors (Invitrogen, USA) and a minimum of six bacterial colonies for each IP_3_R gene were sequenced using M13 forward and reverse primers (Azenta, NJ, USA). To determine the distribution of mutant alleles, sequences were mapped to the genomic DNA using SnapGene.

**Table 2.**
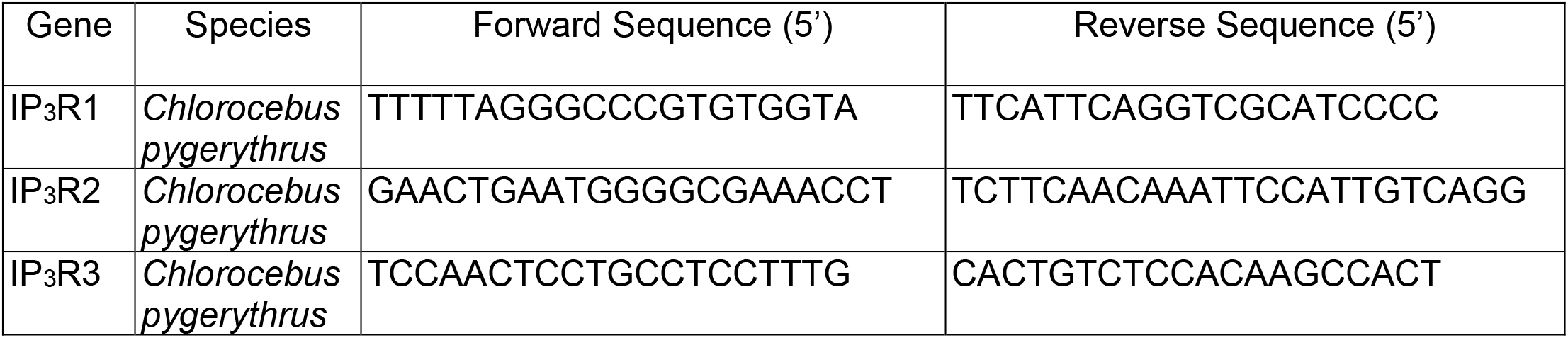
African green monkey IP_3_R1, IP_3_R2, and IP_3_R3 sequencing primers.

### LLC-MK2-GCaMP6s P2Y1 knock-in generation

LLC-MK2-GCaMP6s P2Y1 knock in lines were generated by lentiviral transduction of the human P2RY1 receptor. The P2Y1 cDNA clone was purchased from GenScript (Piscataway, NJ, USA), subcloned into pLVX-IRES-Neo (Takara Bio), and packaged by the BCM Vector Development Core. LLC-MK2-GCaMP6s cells were transduced with MOI 10 in 10 μg/mL polybrene and at 72 hr post-transduction selected using 500 μg/mL G418. (21). Presence of functional P2Y1 was verified by Ca^2+^ imaging for a Ca^2+^ response to 25 nM ADP.

### Ca^2+^ agonist treatment

MA104-GCaMP6s and MA104-GCaMP6s-IP_3_R-TKO monolayers were placed in Ca^2+^ free Hank’s Balanced Salt Solution (HBSS, Invitrogen) and incubated for ∼1 hr to equilibrate. For agonist testing, we used 50 µM ADP for purinergic receptors, 0.75 µM AC55541 or 2 µM Worthington’s trypsin for PAR2 receptors, or 1 µM thapsigargin to inhibit SERCA pumps. Cells were first imaged for 30 seconds to establish a baseline. Following agonist stimulation with 50 µL of agonist solution, cells were imaged for 2 minutes to capture the peak Ca^2+^ response. Imaging acquisition was 50 ms exposures with a 1 second interval.

### Live-cell Ca^2+^ imaging of virus infection

Cell monolayers were grown in Ibidi 8-well chamber slides. On the day of imaging, media was replaced with FluoroBrite-Plus media [FluoroBrite DMEM supplemented with 1X nonessential amino acids, 1X Glutamax, 1X sodium pyruvate, and 15 μM HEPES (Invitrogen)]. Rotavirus was diluted in FluoroBrite-Plus and cells were mock-(FluoroBrite-Plus alone) or rotavirus-infected at MOI 0.01. The inoculum was removed after 1 hr, fresh FluoroBrite-Plus was added, and the slide was mounted into the microscope environmental chamber (37°C in 5% CO_2_) to equilibrate. Imaging was performed using a Nikon TiE inverted microscope using a SPECTRAX LED light source (Lumencor) and either a 20x Plan Fluor (NA 0.45) or a 20X Plan Apo (NA 0.75), objective as previously described (20). Nikon Elements Advanced Research v4.5 software was used for data acquisition and image analysis. Fluorescence intensity values were exported to Microsoft Excel and GraphPad Prism for data analyses.

### Viral infectivity assay

MA104-GCaMP6s and MA104-GCaMP6s-IP_3_R-TKO monolayers grown in 96-well plates were inoculated with 2-fold serial dilutions of SA11-mRuby. The inoculum was removed 1 hr post-infection, then the cells were rinsed twice with PBS and cultured in FBS-free DMEM for ∼16 hrs. Monolayers were fixed in ice cold methanol for 20 minutes at 4°C, washed three times with PBS, and immunostained for 2 hrs using a rabbit anti-rotavirus antisera (strain Alabama) at 1:1000 in PBS, followed by incubation for 1 hr with goat anti-rabbit IgG:AlexaFluor555 at 1:1000 in PBS, and quantitation by fluorescence microscopy.

### Single-cycle and multi-cycle rotavirus yield assay

MA104-GCaMP6s and MA104-GCaMP6s-IP_3_R-TKO monolayers were inoculated with SA11- mRuby for 1 hr. We used an MOI of 0.01 for single-cycle replication assays and 24 PFU/well for multi-cycle replication assays. For single-cycle assays, cells were maintained in DMEM without trypsin, which prevents rotavirus spread due to lack of VP4 spike protein cleavage and harvested at 24 hpi. For multi-cycle assays, cells were maintained in DMEM with 1 µg/mL Worthington’s trypsin and harvested at 24-, 48-, 72-, and 96-hpi. Virus titration was performed by plaque assay (see below).

### Plaque assay

Monolayers of MA104 or LLC-MK2 cells, or their modified derivatives, were inoculated with 10- fold serial dilutions of virus samples in FBS-free DMEM. After the inoculum was removed, 3 mL of overlay (1.2% Avicel in FBS-free DMEM supplemented with 0.1 mg/mL DEAE dextran and 1 μg/mL Worthington’s trypsin) was added to each well and plates were incubated for 3-4 days. After removing the overlay, plates were washed and stained with crystal violet or imaged by fluorescence microscopy prior to crystal violet staining.

### Immunoblot analysis

Confluent MA104-GCaMP6s and MA104-GCaMP6s-IP_3_R-TKO cells inoculated with SA11- mRuby (MOI 10) were harvested in RIPA buffer at 2, 4, 6, 8, and 10 hpi (20). Samples were boiled for 10 minutes at 100°C in SDS-PAGE sample buffer, separated on Tris-glycine 4–20% SDS-PAGE gels (BioRad), and blotted to nitrocellulose, which was blocked with 10% non-fat dry milk in PBS (BLOTTO). Antibodies were diluted in 0.1% BLOTTO and were: rabbit anti-rotavirus [strain Alabama, (1:1000)] (20), rabbit antisera to SA11 NSP4-aa120-146 (1:2000) (24), and mouse anti-GAPDH (1:2000).

### Immunofluorescence

MA104 cell monolayers infected with SA11-mRuby (MOI 1) were fixed at 9 hpi, washed, and incubated overnight at 4°C with rabbit antisera against NSP4 (anti-NSP4-aa114-135) and guinea pig antisera against NSP5 (gift from Dr. Mary Estes at Baylor College of Medicine)(24, 25). Monolayers were washed and incubated for 2 hours at room temperature with fluorescent-conjugated secondary antibodies [anti-Rabbit DyLight 488; anti-guinea pig DyLight 549 (Rockland)] diluted 1:2000 in PBS. Monolayers were imaged using a 63X objective on a Zeiss LSM980 confocal microscope with Airyscan. Mander’s coefficients were determined by calculating the ratio of signal intensity for NSP4 colocalized with that of NSP5 to the overall intensity of the NSP4 signal in a given image following background subtraction. Analyses used three images per group for a total of 46 MA104 parental and 63 MA104-IP_3_R-TKO cells.

### Detection of ADP release from cells

Confluent MA104-GCaMP6s and MA104-GCaMP6s-IP_3_R-TKO cells were infected with SA11- mRuby (MOI 10) and then maintained in 300 µL FluoroBrite-Plus with 20 µM ARL67156, an ecto-ATPase inhibitor. At ∼5 hpi, the supernatant was removed and used to treat parental MA104-GCaMP6s cell monolayers to evoke a Ca^2+^ response, which was monitored by live imaging of GCaMP6s fluorescence and quantitated by measuring the maximum increase in GCaMP6s fluorescence after addition of the supernatant. Specificity of the Ca^2+^ response for P2Y1 activation was determined by pretreating cells with 10 µM BPTU, a P2Y1-selective blocker.

### Quantitative PCR

LLC-MK2-GCaMP6s and LLC-MK2-GCaMP6s+P2Y1 cells were grown to confluency in a 24- well plate. Total RNA was extracted from cell monolayers using TRIzol (Ambion) according to the manufacturer’s instructions. 300 ng of RNA was used to generate cDNA with SensiFAST synthesis reagents (Bioline). Quantitative PCR was performed with Fast SYBR green (Applied Biosystems) with a QuantStudio 3 thermocycler. Purinergic receptor genes were normalized to 18s and expression relative to the lowest expressed gene (P2Y12) was calculated using the ΔΔC_t_ method. Primer sequences have been published previously and can be found in Table 3 (26).

**Table 3.**
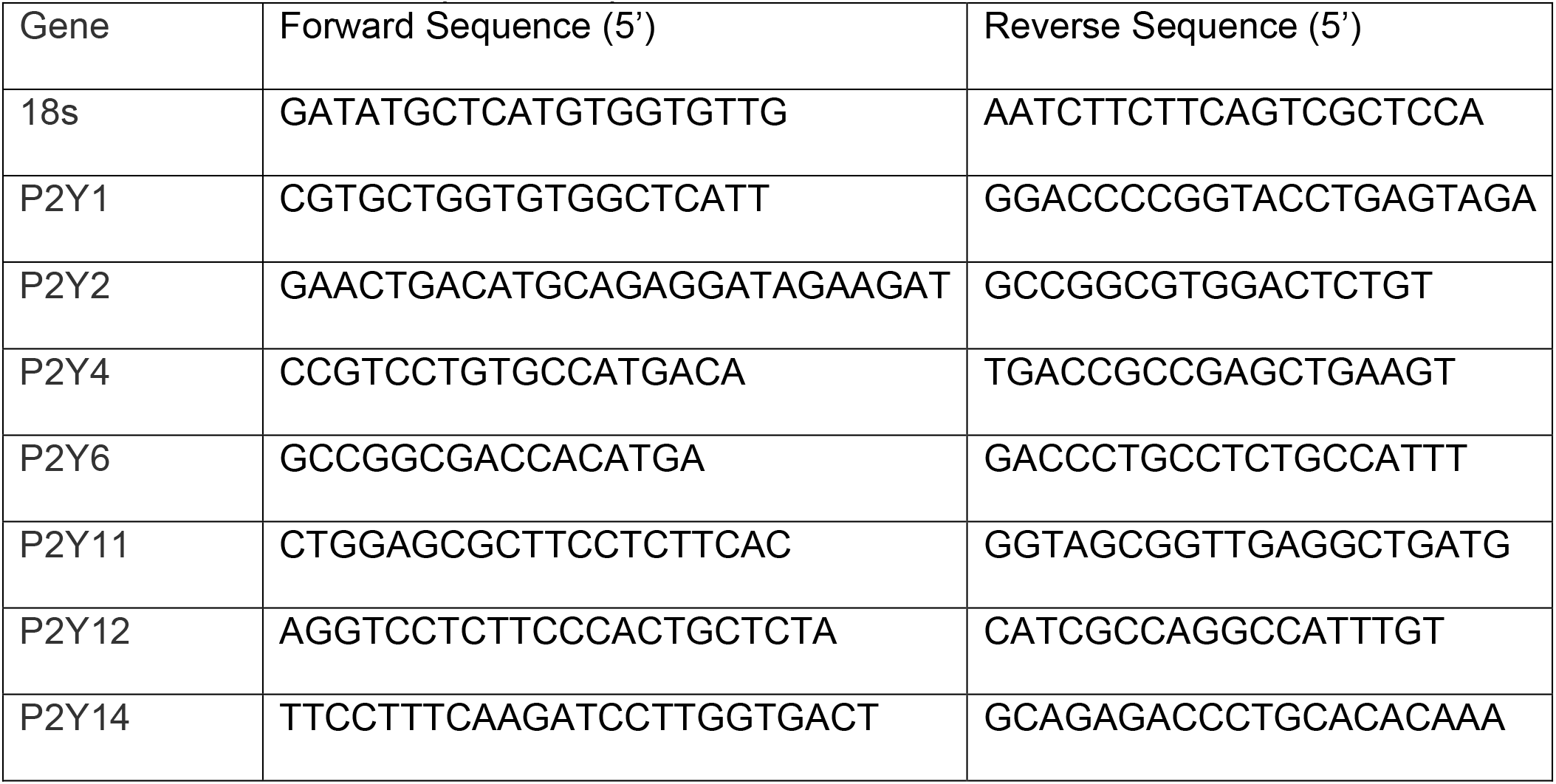
Quantitative PCR primer sequences.

### SA11-mRuby detection in plaques

Rotavirus SA11-mRuby plaques were visualized by fluorescence microscopy using a Nikon TiE inverted microscope as described above. Images were taken from a 6-well tissue culture plate with a 10X Plan Fluor objective (NA 0.30). Whole-well images were obtained by taking a 15 mm x 15 mm stitch with blending and 10% overlap.

### Statistical analyses

Statistics were completed using GraphPad Prism (version 8.4.3). Unless stated, all experiments were performed in biological triplicate. The threshold for Ca^2+^ spikes is a 5% increase in GCaMP fluorescence, as previously defined for this system (20). We performed column statistics to determine the normality of the data sets. We used unpaired Student’s t test for data sets with a parametric distribution or a Mann-Whitney test for data sets with a nonparametric distribution. We applied a one-way ANOVA with proper correction when applicable for comparing sample groups to control groups. Differences were determined statistically significant if the P value was <0.05. For all figures, P value notations are as follows: *P <0.05, **P <0.01, ***P <0.001, and ****P <0.0001.

## Results

Previous studies have shown that rotavirus NSP4’s viroporin function is a critical mechanism for the dysregulation of cellular Ca^2+^ homeostasis during infection (7, 20). Yet, NSP4 crosstalk with host IP_3_R channels could contribute to the overall Ca^2+^ signaling in infected cells and the recently discovered intercellular Ca^2+^ waves (ICWs) induced by rotavirus are generated by IP_3_R Ca^2+^ release upon activation of P2Y1 receptors (14-17). Thus, we investigated the role of IP_3_R in the rotavirus-induced Ca^2+^ signaling landscape using IP_3_R-null cell lines generated by CRISPR-Cas9 genome editing. First, we obtained IP_3_R triple-knockout (IP_3_R-TKO) HEK293 cells, which express no functional IP_3_R isoforms, and tested whether rotavirus infection elevates Ca^2+^ signaling similar to MA104 cells (15). Using long-term, time-lapse Ca^2+^ imaging, we found that rotavirus infected MA104-GCaMP6s cells exhibited increased Ca^2+^ signaling, as we previously described (20). Representative images show both the low amplitude intracellular Ca^2+^ signals observed in rotavirus-infected cells (at 354 min and 360 min), as well as high amplitude ICWs that propagate to surrounding uninfected cells (396 min) (Figure 1A, Movie S1). Representative traces show mock-infected cells exhibit no strong Ca^2+^ signaling events (Figure 1B, black line), but rotavirus-infected cells exhibited a strong increase in dynamic Ca^2+^ signaling (Figure 1B, red line). Further, the intracellular Ca^2+^ signals, highlighted in the magenta bracket, occur before the onset of ICWs (Figure 1B). This increase in Ca^2+^ signaling results in significantly more Ca^2+^ spikes in rotavirus-infected than in mock infected cells (Figure 1C).

**Figure 1.**
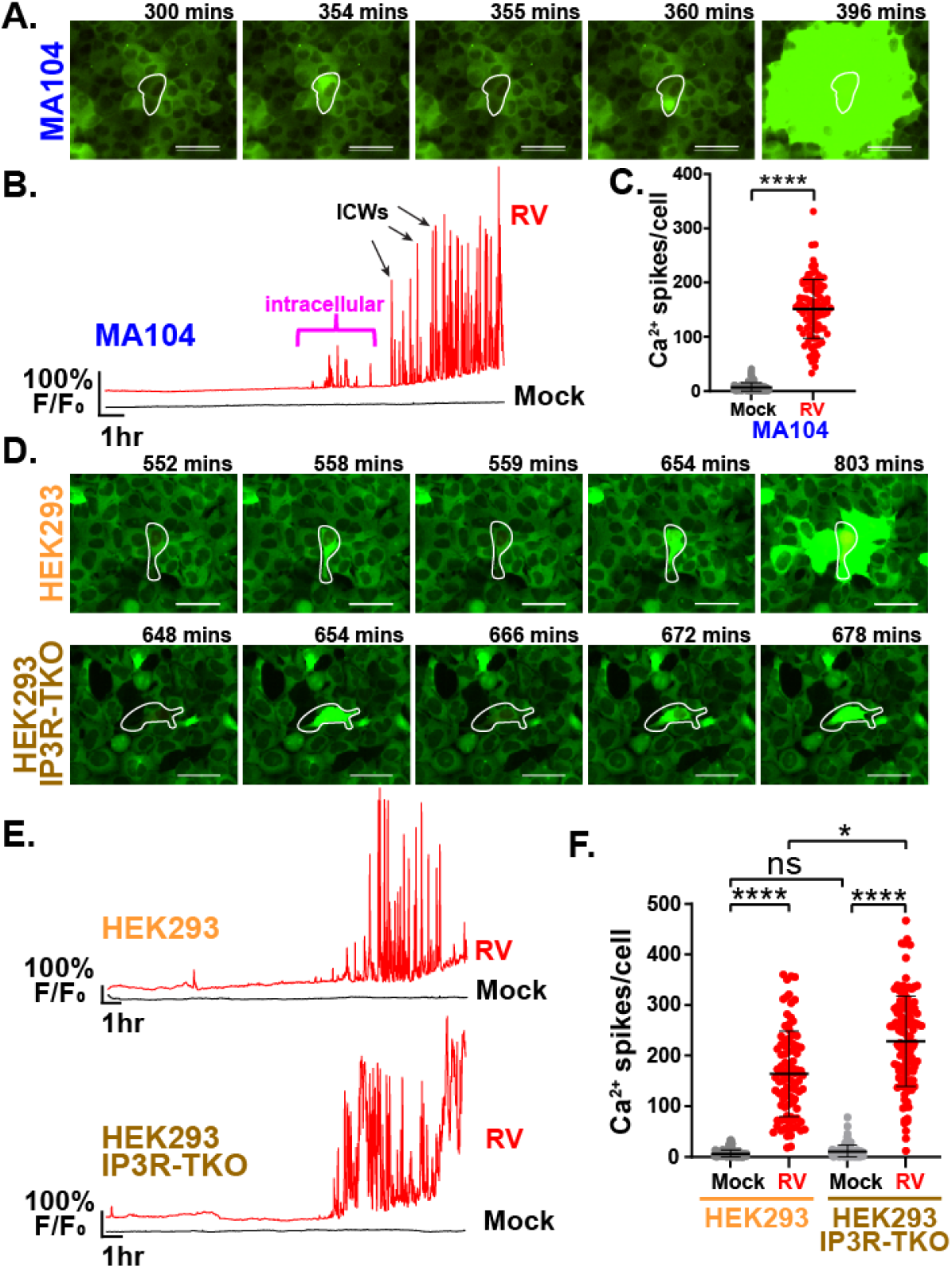
Time lapse Ca^2+^ imaging of rotavirus (RV) infected MA104-GCaMP6s, HEK293- GCaMP6s, and HEK293-GCaMP6s-IP_3_R-TKO cells. **A.** Filmstrip of a RV infected MA104 cell (white outline) showing both intracellular and intercellular Ca^2+^ signals. **B.** Normalized GCaMP6s fluorescence (F/F_0_) of representative mock (black) and RV-infected (red) cells from ∼2-20 hpi. Intracellular Ca^2+^ signals (magenta bracket) occurred prior to the onset of ICWs (black arrows = representative ICWs). **C.** Quantitation of Ca^2+^ spikes per cell in mock (grey) or RV-infected (red) cells. ****p<0.001 by Mann-Whitney T test. **D.** Filmstrip images of representative HEK293- GCaMP6s and HEK293-GCaMP6s-IP_3_R-TKO cells (white outline) showing intracellular Ca^2+^ signals over time. **E.** Representative single-cell of normalized GCaMP6s fluorescence from mock (black) and RV-infected (red) HEK293-GCaMP6s and HEK293-GCaMP6s-IP_3_R-TKO cells. **F.** Quantitation of Ca^2+^ spikes per cell in mock (grey) or RV-infected (red) HEK293- GCaMP6s and HEK293-GCaMP6s-IP_3_R-TKO cells. Statistical analysis by Kruskal-Wallis with Dunn’s multiple correction where *p<0.05 and ****p<0.0001. All data are displayed as mean ± SD. Scale bars are 50 µm.

Next, to test the role of IP_3_R in dysregulation of Ca^2+^ signaling during rotavirus infection, we examined Ca^2+^ signaling in SA11-mRuby infected parental HEK293-GCaMP6s and HEK293- GCaMP6s-IP_3_R-TKO cells (Figure 1D-F). Representative images (Figure 1D) and Ca^2+^ signaling traces (Figure 1E) show that both parental HEK293-GCaMP6s and HEK293-GCaMP6s-IP_3_R-TKO cells exhibit robust Ca^2+^ signaling during rotavirus infection (red lines) that is much greater than in mock-infected cells (black lines). We observe numerous intracellular Ca^2+^ signals in both cell lines but only a few, small ICWs in parental HEK293-GCaMP6s cells. There were no ICWs in the HEK293- GCaMP6s-IP_3_R-TKO cells because, as a Gq-coupled GPCR, P2Y1 evokes an IP_3_R-mediated release of ER Ca^2+^ (Figure 1D and Movie S2). Quantitation of Ca^2+^ signaling in parental HEK293-GCaMP6s and HEK-293-GCaMP6s-IP_3_R-TKO cells show both cell types had a significant increase in overall spikes per cell compared to mock cells (Figure 1F). While we observed a small, but statistically significant, increase in overall Ca^2+^ signaling in HEK293- GCaMP6s-IP_3_R-TKO cells compared to parental HEK293-GCaMP6s cells (Figure 1F), the critical observation is that there was no *decrease* in rotavirus-induced Ca^2+^ signaling in HEK293-GCaMP6s-IP_3_R-TKO cells. Together, these data show rotavirus infection produces robust Ca^2+^ signaling in both MA104 and HEK293 cells, and this is not decreased in the absence of IP_3_R. However, HEK293 cells are suboptimal for examining rotavirus-induced Ca^2+^ signaling, since they do not exhibit robust ICWs and therefore do not have distinct intracellular and intercellular signaling patterns. Thus, for further investigation into the role of IP_3_R-mediated signaling during rotavirus infection, we aimed to establish an MA104-GCaMP6s-IP_3_R-TKO cell line.

### Generation MA104-GCaMP6s cells lacking IP_3_R expression

To generate an MA104-GCaMP6s cell line lacking IP_3_R expression, we generated a CRISPR-Cas9 lentivirus construct that expresses individual gRNAs targeted to each IP_3_R gene (Table 1). Our workflow to generate MA104-GCaMP6s-IP_3_R-TKO cells is depicted in Figure 2A. For screening and validation, we used two different GPCR agonists to activate IP_3_R-mediated Ca^2+^ responses: 50 µM ADP to activate P2Y purinergic receptors and 0.75 µM AC55541 to activate the Protease-activated Receptor 2 (PAR2). We identified 8 clones that exhibited little to no response to these two agonists (Figure 2B) and chose clone F3 for full characterization, which will be designated MA104-GCaMP6s-IP_3_R-TKO cells for the remainder of this paper.

**Figure 2.**
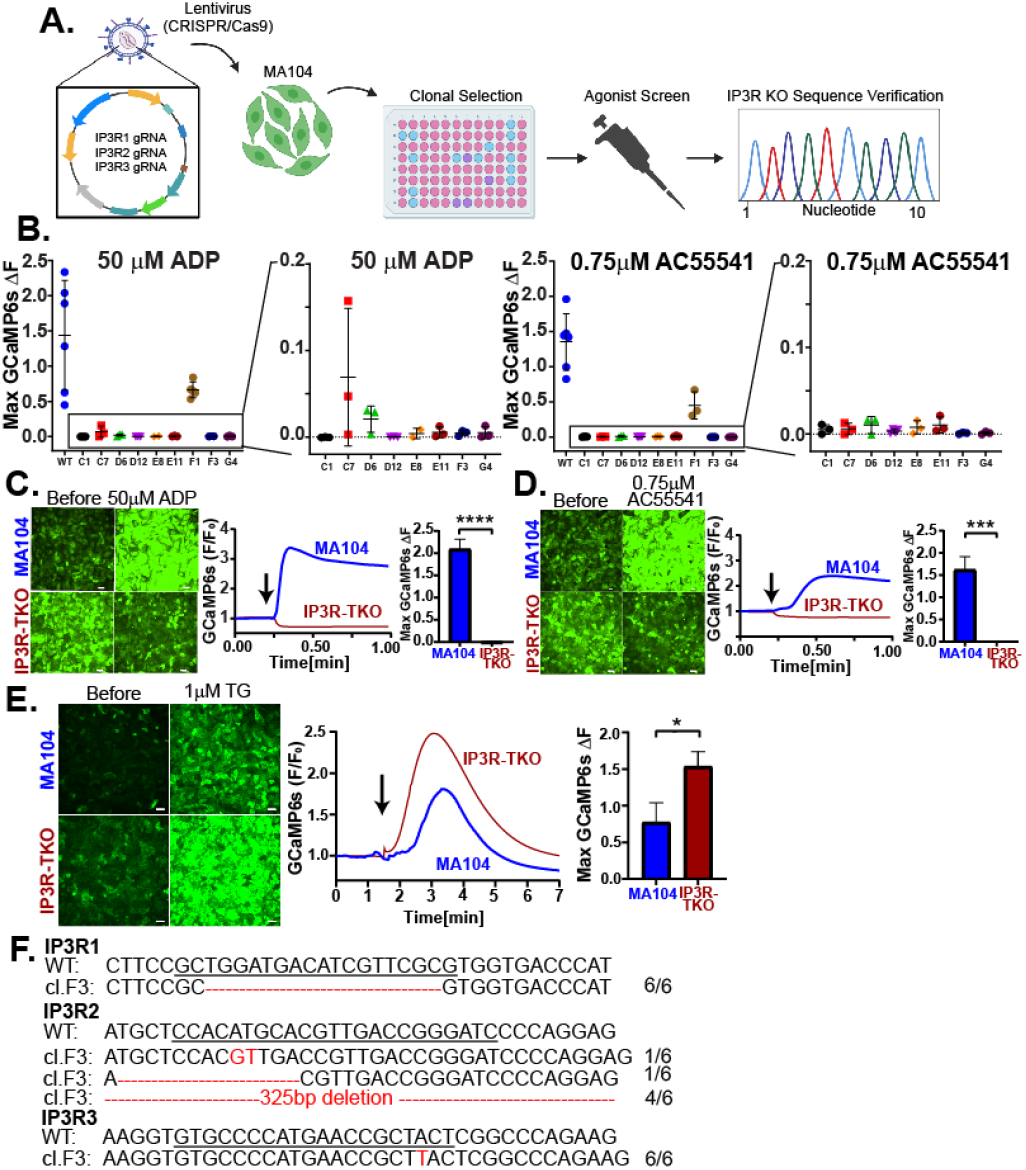
Development of MA104-IP_3_R-TKO GCaMP6s cell line. **A.** Schematic workflow for the generation and validation of the MA104-GCaMP6s-IP_3_R-TKO cells. **B.** Maximum change in GCaMP6s fluorescence in MA104-GCaMP6s and MA104-GCaMP6s-IP_3_R-TKO clones after agonist screening using ADP and AC55541 (PAR2 agonist). **C.** (Left) MA104-GCaMP6s and MA104-GCaMP6s-IP_3_R-TKO cells before and after treatment with ADP. Traces (middle) and quantitation (right) of GCaMP6s fluorescence showing Ca^2+^ responses of parental and IP_3_R-KO cells upon ADP addition (arrow). **D.** Agonist treatment studies as in (**C**) but using 0.75 µM AC55541. **E.** MA104-GCaMP6s and MA104-GCaMP6s-IP_3_R-TKO cells before and after 1 µM thapsigargin treatment. Ca^2+^ signaling traces and quantitation showing normalized GCaMP6s fluorescence after 1 µM thapsigargin treatment (arrow). **F.** Genomic DNA sequencing of IP_3_R1, IP_3_R2, and IP_3_R3 gRNA sites for MA104-GCaMP6s and MA104-GCaMP6s-IP_3_R-TKO cells. All data represented as mean ± SD. Statistical analyses were performed by Mann-Whitney test with *p<0.05, ***p<0.001, ****p<0.0001. Scale bars are 50 µm.

To functionally validate the loss of IP_3_R Ca^2+^ signals in MA104-GCaMP6s-IP_3_R-TKO cells, we performed live imaging to measure Ca^2+^ responses to ADP and PAR2 agonist AC55541 (Figure 2C-D). For both agonists, MA104-GCaMP6s-IP_3_R-TKO cells exhibit no increase in Ca^2+^ and a small, but reproducible, decrease in GCaMP6s fluorescence (Figure 2C-D, red traces; Movie S3). To validate these cells, maintain an ER Ca^2+^ store, we treated cells with thapsigargin to inhibit SERCA pumps. We found thapsigargin induced an increase in GCaMP6s fluorescence for both parental and IP_3_R-TKO cells, and peak signal from IP_3_R-TKO cells was significantly greater than that of parental MA104-GCaMP6s cells (Figure 2E). This indicates that IP_3_R-TKO cells have a higher ER Ca^2+^ load than that of parental MA104-GCaMP6s cells, which would be expected in the absence of basal IP_3_R activity. Finally, sequence analyses of MA104-GCaMP6s-IP_3_R-TKO cells identified the mutations at the gRNA sites for each IP_3_R gene. To assess the distribution of mutations, we amplified the gRNA sites for each gene, PCR-cloned amplicons, and sequenced 6 plasmids for each IP_3_R gene. We found a deletion mutation for IP_3_R1, three different insertion/deletion mutations for IP_3_R2 and a single allele insertion for IP_3_R3 (Figure 2F), but no wild-type sequences were isolated, consistent with the lack of agonist-induced Ca^2+^ responses from these cells. Together, these data show that we have successfully generated an MA104-GCaMP6s cell line lacking IP_3_R expression to better examine the role of IP_3_R-mediated ER Ca^2+^ release during rotavirus infection.

### Rotavirus-induced Ca^2+^ signaling in the absence of IP_3_R

We previously found that rotavirus infection results in activation of highly dynamic Ca^2+^ signaling throughout infection (20). Now by generating MA104-GCaMP6s-IP_3_R-TKO cells, we set out to determine how the lack of IP_3_R affects rotavirus-induced Ca^2+^ signaling. We performed long-term, time-lapse Ca^2+^ imaging of both parental MA104-GCaMP6s and MA104-GCaMP6s-IP_3_R-TKO cells infected with SA11-mRuby rotavirus, and representative images are shown in Figure 3A. As previously characterized, rotavirus infected parental MA104-GCaMP6s cells exhibit an increase in intracellular Ca^2+^ signals present, as well as production of intercellular Ca^2+^ waves (ICWs) (Figure 3A, top; Movie S4). In MA104-GCaMP6s-IP_3_R-TKO cells, we still observed robust intracellular Ca^2+^ signaling in rotavirus-infected cells, but no ICWs were produced (Figure 3A, bottom; Movie S4). Next, we compared Ca^2+^ signaling phenotypes between parental MA104- GCaMP6s and MA104-GCaMP6s-IP_3_R-TKO cells, with 4 representative traces for each shown in Figure 3B. We found both cell types exhibit a very similar pattern of robust Ca^2+^ signaling that progressively increases over the course of infection but mock-infected cells exhibited only basal level signaling (Figure 3B). Further, though both cell lines show increased Ca^2+^ signaling dynamics, there were fewer high amplitude Ca^2+^ signals in MA104- GCaMP6s-IP_3_R-TKO cells, which likely represents autocrine activation of P2Y1 during ICWs (Figure 3B, arrows). Next, we quantitated the number of Ca^2+^ spikes in rotavirus-infected and uninfected neighboring cells 3 and 5 cells away (NB3 and NB5, respectively) to capture ICWs (Figure 3C). In parental MA104-GCaMP6s cells, we found a significant increase in Ca^2+^ signaling in both infected and neighboring cells, consistent with our previous studies (20). In MA104-GCaMP6s-IP_3_R-TKO cells, there was a significant increase in Ca^2+^ signaling in rotavirus-infected cells but no increase in the neighboring cells, consistent with the lack of ICWs (Figure 3C). Interestingly, in rotavirus-infected MA104-GCaMP6s-IP_3_R-TKO cells, we found no significant decrease in overall Ca^2+^ signaling (Figure 3C). Since MA104-GCaMP6s-IP_3_R-TKO cells did not produce ICWs, we wanted to determine whether the cells were still releasing ADP during infection. To test this, parental MA104-GCaMP6s and MA104-GCaMP6s-IP_3_R-TKO cells were mock or rotavirus-infected and incubated in the presence of ARL67156, an ecto-nucleotidase inhibitor, to stabilize extracellular purines. At 5 hpi, a time point when ICWs have begun, media was removed from mock and infected cells and added to parental MA104- GCaMP6s cells. We found that media from rotavirus-infected cells induced a significantly greater Ca^2+^ response than that from mock infected cells, and the Ca^2+^ response was blocked by the P2Y1 inhibitor BPTU (Figure 3D). Together, this indicated that the loss of IP_3_R does not reduce the rotavirus-induced release of ADP during infection and that the IP_3_R-independent arms of the P2Y1 signaling pathway are likely intact. These data in MA104 cells echo that of our initial studies in HEK293-GCaMP6s-IP_3_R-TKO cells, in that rotavirus still dramatically increases Ca^2+^ signaling within infected cells in the absence of IP_3_R (Figure 1E). Yet, loss of IP_3_R resulted in the abrogation of rotavirus induced ICWs and significantly reduced the increase in Ca^2+^ signaling in surrounding uninfected cells, which ultimately changes the overall Ca^2+^ signaling landscape during infection.

**Figure 3.**
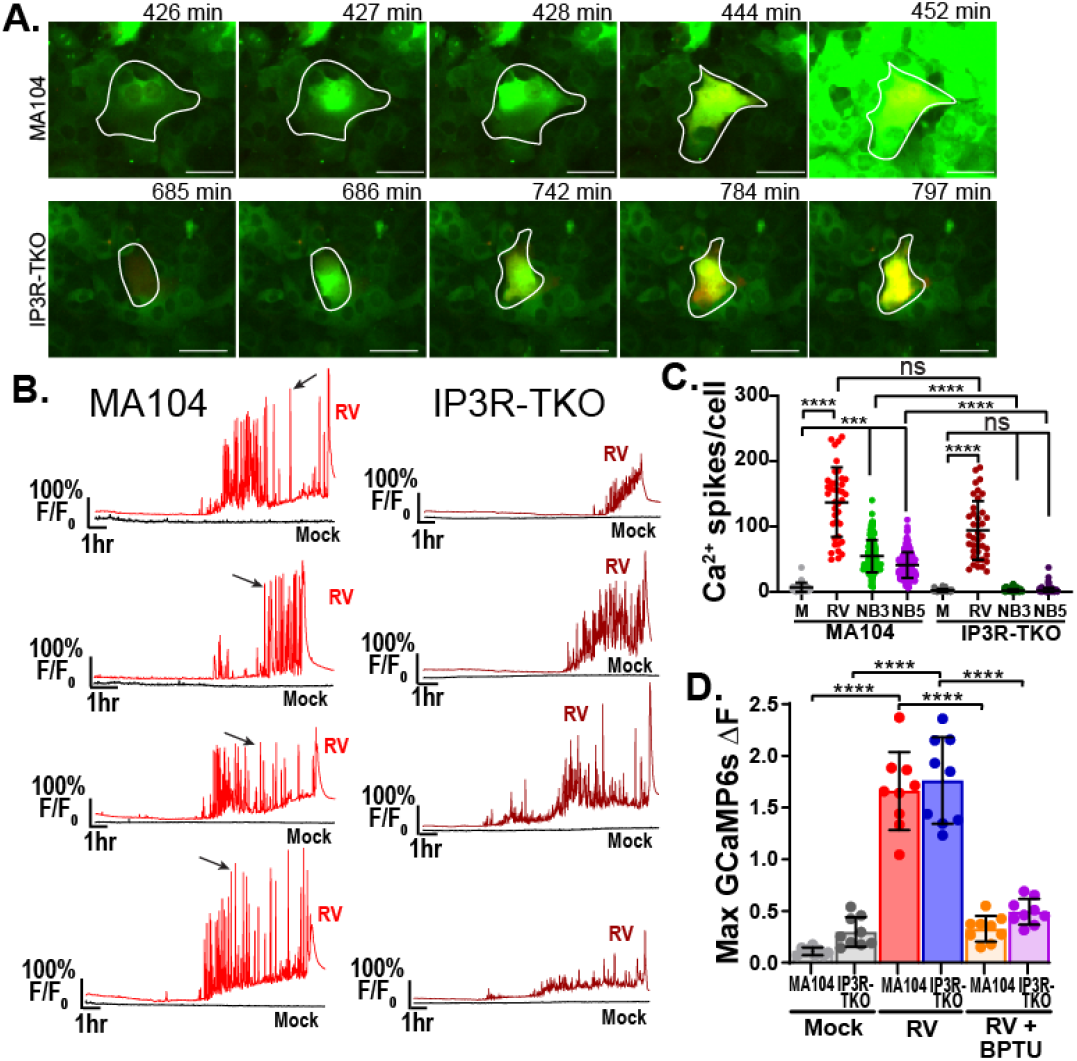
Characterization of MA104-GCaMP6s and MA104-GCaMP6s-IP_3_R-TKO cells (IP_3_R-TKO) during RV infection. **A.** Filmstrip images of RV infected MA104-GCaMP6s cells (white outline, top row) and MA104-GCaMP6s-IP_3_R-TKO cells (white outline, bottom row) overtime. **B.** Four representative single-cell traces of normalized GCaMP6s fluorescence (F/F_0_) from mock (black) or RV-infected MA104-GCaMP6s cells (red) or MA104-GCaMP6s-IP_3_R-TKO cells (maroon) Arrows indicate example occurrences of an intercellular Ca^2+^ wave signal. **C.** Quantitation of the number of Ca^2+^ spikes per cell in mock (grey), RV-infected and neighboring cells for MA104-GCaMP6s and MA104-GCaMP6s-IP_3_R-TKO cells. **D.** Maximum change in normalized GCaMP6s fluorescence after supernatant transfer of RV infected MA104-GCaMP6s (red) and MA104-GCaMP6s-IP_3_R-TKO (blue) cells to MA104-GCaMP6s sensor cells treated with 10µM BPTU. All data are shown as the mean ± SD. Statistical analyses were performed using Kruskal-Wallis with Dunn’s multiple corrections tests with ***p<0.001 & ****p<0.0001. Scale bars are 50 µm.

### Role of IP_3_R in rotavirus infection and replication

Numerous studies have confirmed that increased cytosolic Ca^2+^ is critical for rotavirus replication (12, 13, 27). Thus, we next used parental MA104-GCaMP6s and MA104-GCaMP6s-IP_3_R-TKO cells to assess whether IP_3_R signaling plays a role in rotavirus infectivity and/or replication. First, we compared SA11-mRuby plaque phenotypes between parental MA104- GCaMP6s and MA104-GCaMP6s-IP_3_R-TKO cells, discovering a major difference in plaque morphology (Figure 4A). MA104-GCaMP6s cell plaques fully cleared the monolayer and had clean margins (Figure 4A, Top) in contrast to MA104-GCaMP6s-IP_3_R-TKO cells, which exhibited a similar number of plaques (i.e., no difference in measured titer); however, the plaques were turbid with most of the monolayer remaining intact (Figure 4A, Bottom). Quantification of plaque diameters showed rotavirus formed significantly smaller plaques on MA104-GCaMP6s-IP_3_R-TKO cells than on parental MA104- GCaMP6s cells (Figure 4A). Due to this substantial difference in plaque formation, we next determined whether loss of IP_3_R affected rotavirus infectivity or single-cycle virus yield. We first determined infectivity of SA11-mRuby for both parental MA104-GCaMP6s and MA104- GCaMP6s-IP_3_R-TKO cells using a fluorescent focus assay (FFA) and found no differences in the observed titer, indicating that rotavirus infects both cell lines with the same efficiency (Figure 4B). This is consistent with both our live imaging studies (Figure 3) and plaque assays (Figure 4A), in which we observed a similar number of infected cells and plaques, respectively. Next, we tested whether the MA104- GCaMP6s-IP_3_R-TKO cells efficiently supported rotavirus replication. We compared single-cycle virus yield between parental MA104-GCaMP6s and MA104-GCaMP6s-IP_3_R-TKO cells. Cells were infected with MOI 0.01 and maintained in the absence of trypsin to limit the infection to a single round. We found that yields were similar between parental MA104-GCaMP6s and MA104-GCaMP6s-IP_3_R-TKO cells, though there was a small increase in rotavirus yield in MA104- GCaMP6s-IP_3_R-TKO cells (Figure 4C). Thus, the small plaque phenotype on the MA104- GCaMP6s-IP_3_R-TKO cells cannot be explained by an inability of these cells to support rotavirus replication. We evaluated rotavirus protein expression by western blot and found similar levels and kinetics of viral protein synthesis for both structural proteins and NSP4 in both cell lines (Figure 4D). Finally, we using immunofluorescence staining of rotavirus-infected cells, we found no difference in viroplasm formation localization of NSP4 to viroplasms (Figure 4E) and this was confirmed by determining no significant difference in the Manders’ coefficient for each cell (MA104: 0.047±0.018; MA104-IP_3_R-TKO: 0.034±0.013, p=0.7). Together, these data indicate the loss of IP_3_R did not significantly affect the ability of rotavirus to infect, produce proteins, or assemble new progeny virus during the initial round of replication, and therefore suggests the defect in plaque formation occurs during rotavirus spread through multiple rounds of replication.

**Figure 4.**
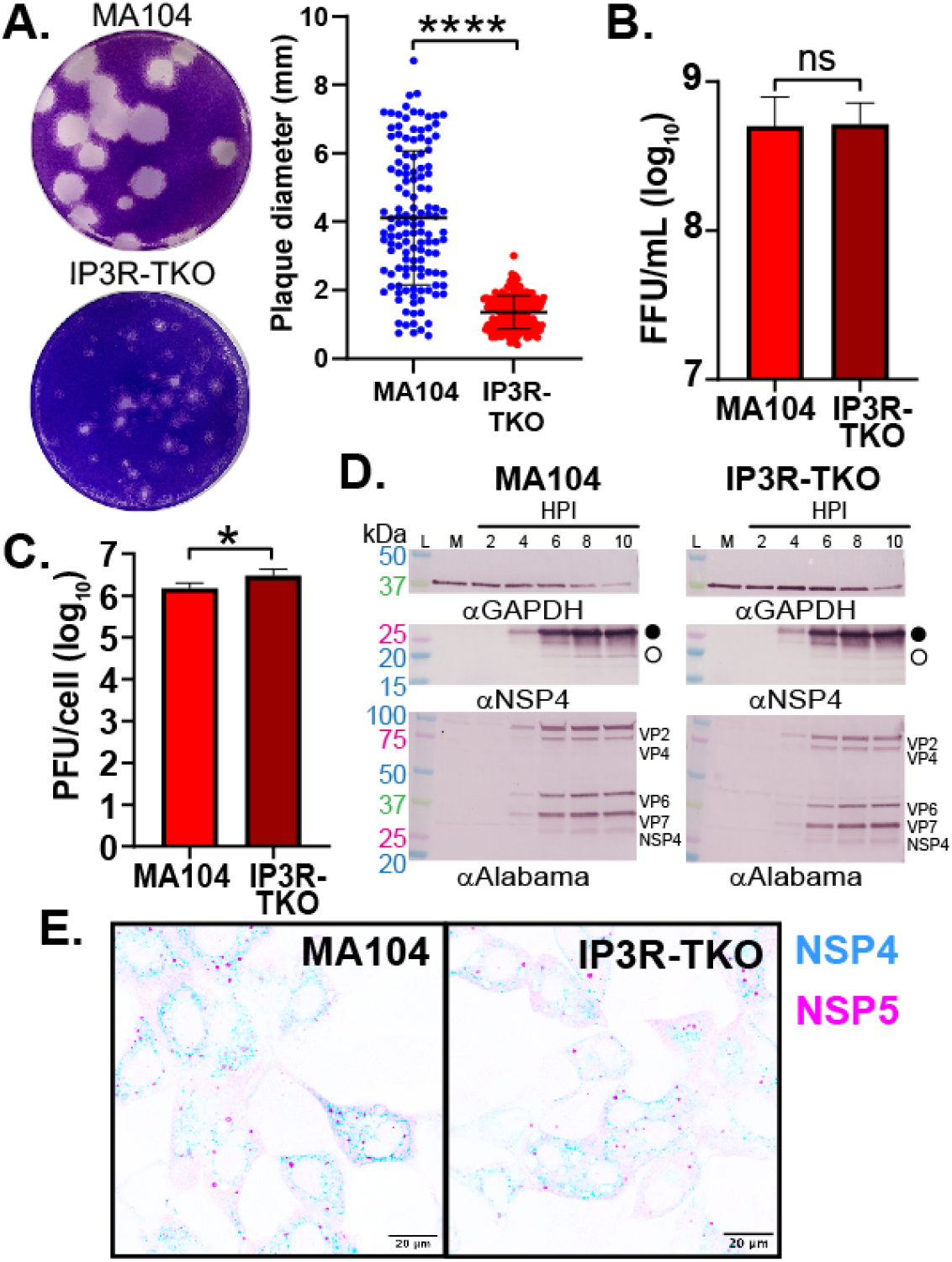
Characterization of rotavirus infectivity and replication in MA104-GCaMP6s-IP_3_R-TKO cells. **A.** Representative images and diameter quantitation of RV plaques formed on MA104- GCaMP6s or MA104-GCaMP6s-IP_3_R-TKO cells. **B.** Quantitation of rotavirus infectivity by FFA on MA104-GCaMP6s (red) or MA104-GCaMP6s-IP_3_R-TKO (dark red) cells. **C.** One-step rotavirus yield from MA104-GCaMP6s (red) or MA104-GCaMP6s-IP_3_R-TKO (dark red) cells was measured by plaque assay and graphed as Log_10_ plaque forming units (PFU) per cell. **D.** Kinetics of rotavirus protein expression in MA104-GCaMP6s and MA104-GCaMP6s-IP_3_R-TKO cells was determined by immunoblot for GAPDH (top), NSP4 (middle), and a rabbit anti-rotavirus (strain Alabama) (bottom). **E.** Immunofluorescence of RV-infected MA104-GCaMP6s and MA104- GCaMP6s-IP_3_R-TKO cells to detect NSP4 (blue) and NSP5 (magenta), as a marker of viroplasms. All data are shown as the mean ± SD. Statistical analyses were performed using Mann-Whitney T tests with *p<0.05 & ****p<0.0001. Scale bars are 20 µm.

### Role of IP_3_R and ICWs in rotavirus spread

To further investigate why rotavirus did not plaque efficiently on MA104-GCaMP6s-IP_3_R-TKO cells, we used fluorescent microscopy to examine the spread of SA11-mRuby in the area around plaques for both parental MA104-GCaMP6s and MA104-GCaMP6s-IP_3_R-TKO cells. We performed plaque assays with SA11-mRuby and, on day 4 post-infection, we replaced the overlay with PBS and imaged plaques by brightfield and fluorescence microscopy to visualize mRuby expression in the plaques. In parental MA104 cells, there was full clearing of the plaque and strong mRuby expression in a wide margin around the plaque (Figure 5A, top). In contrast, plaques on MA104-GCaMP6s-IP_3_R-TKO cells do not clear and the area of mRuby-positive cells were substantially smaller (Figure 5A, bottom).

**Figure 5.**
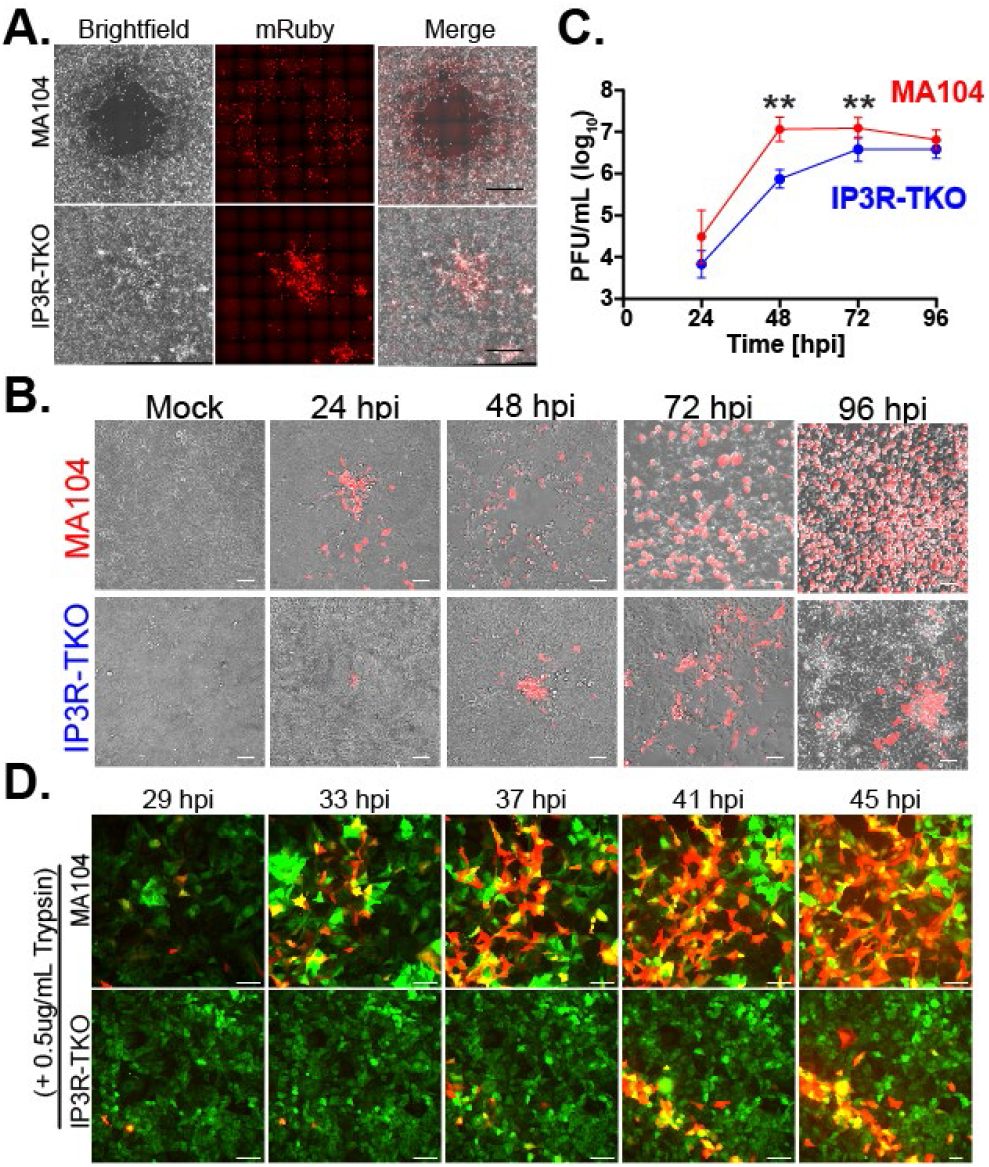
Loss of IP_3_R reduces rotavirus spread. **A.** Visualization of SA11-mRuby in plaques from MA104-GCaMP6s or MA104-GCaMP6s-IP_3_R-TKO cells. Brightfield (left) and SA11-mRuby fluorescence (middle) images are merged on the right. Images were acquired with a Plan Fluor objective (NA 0.30) with 10×10mm stich parameters. Scale bars are 1000 µm **B.** SA11-mRuby spread in MA104-GCaMP6s (red) or MA104-GCaMP6s-IP_3_R-TKO (blue) cells. Brightfield and fluorescence (SA11-mRuby) images were acquired at the indicated times post infection using a 20x Plan Fluor objective (NA 0.45). **C.** SA11-mRuby yield from 24-96 hpi from MA104-GCaMP6s (red) or MA104-GCaMP6s-IP_3_R-TKO cells (blue) was measured by plaque assay and graphed as Log_10_ PFU/mL. **D.** Live imaging of SA11-mRuby (RFP) spread in GCaMP6s-expressing parental MA104-GCaMP6s (top) or MA104-GCaMP6s-IP_3_R-TKO cells (bottom) from ∼30-45 hpi. All data are shown as the mean ± SD. Scale bars are 50 µm in B & D. **p<0.01 by T-test.

Next, we examined whether the smaller, turbid plaques formed on MA104-GCaMP6s-IP_3_R-TKO cells were the result of a decrease in virus replication kinetics in multi-step replication. We infected parental MA104-GCaMP6s and MA104-GCaMP6s-IP_3_R-TKO cells with ∼24 PFU (MOI = ∼5.9×10^-5^) to ensure well isolated infected cells, and cultured in the presence of trypsin to visualize SA11-mRuby spread and measure the kinetics of virus replication over 96-hours post-infection. Using live microscopy, we tracked SA11-mRuby spread throughout the monolayer in both parental MA104-GCaMP6s and MA104-GCaMP6s-IP_3_R-TKO cells (Figure 5B). In parental MA104-GCaMP6s cells, there was more rapid spread with complete destruction of the monolayer by 48-72 hpi (Figure 5B, top). In contrast, virus spread in MA104-GCaMP6s-IP_3_R-TKO cells was substantially delayed, especially between 24-48 hpi (Figure 5B, bottom). Next, we determined virus yield from 24-96 hpi and found parental MA104-GCaMP6s cells supported faster virus replication than MA104-GCaMP6s-IP_3_R-TKO cells, with peak titers reached in parental cells by 48 hpi (Figure 5C, red line). In contrast, virus replication was significantly lower in MA104-GCaMP6s-IP_3_R-TKO cells from 48-72 hpi and did not peak until 72-96 hpi (Figure 5C, blue line).

Finally, we examined rotavirus spread using live, time-lapse microscopy to measure the rate of mRuby expression in the absence or presence of trypsin (Figure 5D). As was shown in Figure 3A, in the absence of trypsin there are single infected cells and no spread of mRuby to neighboring cells (data not shown). In the presence of trypsin, rotavirus spread rapidly from the initial infected cells to many of the surrounding cells in parental MA104- GCaMP6s cells, but in MA104-GCaMP6s-IP_3_R-TKO cells, this was much slower and primarily to adjacent cells (Figure 5D, Movie S5). Together, these data show that in infected cells, the host IP_3_R ER Ca^2+^ channel is largely for dispensable rotavirus-induced in Ca^2+^ signaling; however, IP_3_R-mediated Ca^2+^ signaling was critical for rotavirus replication and spread.

### Priming Uninfected Cells via P2Y1-mediated ICWs

As the loss of IP_3_R did not affect rotavirus infectivity or replication in the single cycle replication studies, the above data indicated the defect in rotavirus spread was related to the lack of increased Ca^2+^ signaling in neighboring cells. Increased Ca^2+^ signaling in rotavirus-infected cells has been shown to be critical for robust replication (12, 28, 29), so these observations led us to hypothesize that the increased Ca^2+^ signaling in neighboring uninfected cells could promote rotavirus replication/spread by priming them for infection. Thus, we set out to test this hypothesis by examining how the manipulation of ICWs affected rotavirus spread.

We previously showed P2Y1 receptor blockers significantly inhibits rotavirus induced ICWs (17). Thus, we next tested whether blocking ICWs with BPTU, a P2Y1 selective blocker, would also reduce rotavirus spread. We performed a plaque assay using parental MA104 cells in the presence of 1 μM BPTU, or DMSO vehicle control, and found BPTU treatment resulted in significantly smaller plaques (Figure 6). Thus, reducing ICWs by blocking P2Y1 causes a similar reduction in the ability of rotavirus to form plaques as the loss of IP_3_R; however, P2Y1 blockers, or even P2Y1 knockout cells, did not fully abrogate ICWs as strongly as MA104-GCaMP6s-IP_3_R-TKO cells (17). Therefore, to further test the role of ICWs in rotavirus replication, we examined rotavirus spread in LLC-MK2-GCaMP6s cells, a rhesus monkey kidney cell line we used previously to study Ca^2+^ signaling by Recoviruses (Rhesus enteric caliciviruses) (22). We observed that while rotavirus-infected LLC-MK2-GCaMP6s cells exhibited increased Ca^2+^ signaling, there were no ICWs and therefore no increase in Ca^2+^ signaling in neighboring uninfected cells (Figure 7A). Further, treatment of LLC-MK2-GCaMP6s cells with ADP did not evoke a Ca^2+^ response (Figure 7B, D), indicating a lack of functional P2Y1 signaling. So, we reasoned that if ICWs are a critical factor in rotavirus spread, exogenous expression of P2Y1 in LLC-MK2-GCaMP6s cells might rescue the rotavirus-induced ICW phenotype and increased rotavirus spread. We used lentivirus transduction to generate a stable P2Y1 knock-in LLC-MK2-GCaMP6s cell line (LLC-MK2-GCaMP6s+P2Y1) and presence of an ADP-stimulated Ca^2+^ signal confirmed the P2Y1 was functional in these cells (Figure 7C-D, Movie S6). We confirmed a significant increase in P2Y1 expression in the LLC-MK2-GCaMP6s+P2Y1 cells by qRT-PCR and found that the P2Y1 knock-in did not affect the expression of any other P2Y purinergic receptor (Figure 7E).

**Figure 6.**
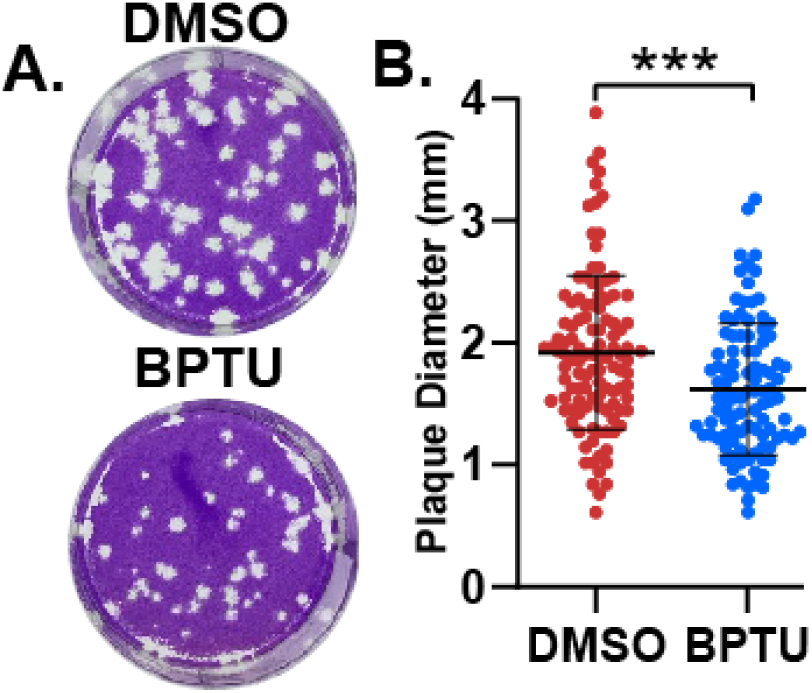
Blocking P2Y1-mediated ICWs reduces rotavirus plaque size. **A.** Representative image of RV plaques formed on parental MA104 cell monolayers treated with DMSO (Top) or 1 µM BPTU (bottom). **B.** Measurement of plaque diameter for DMSO and 1 µM BPTU treated cells. Data are shown as the mean ± SD. ***p<0.001 by Mann-Whitney T test.

**Figure 7.**
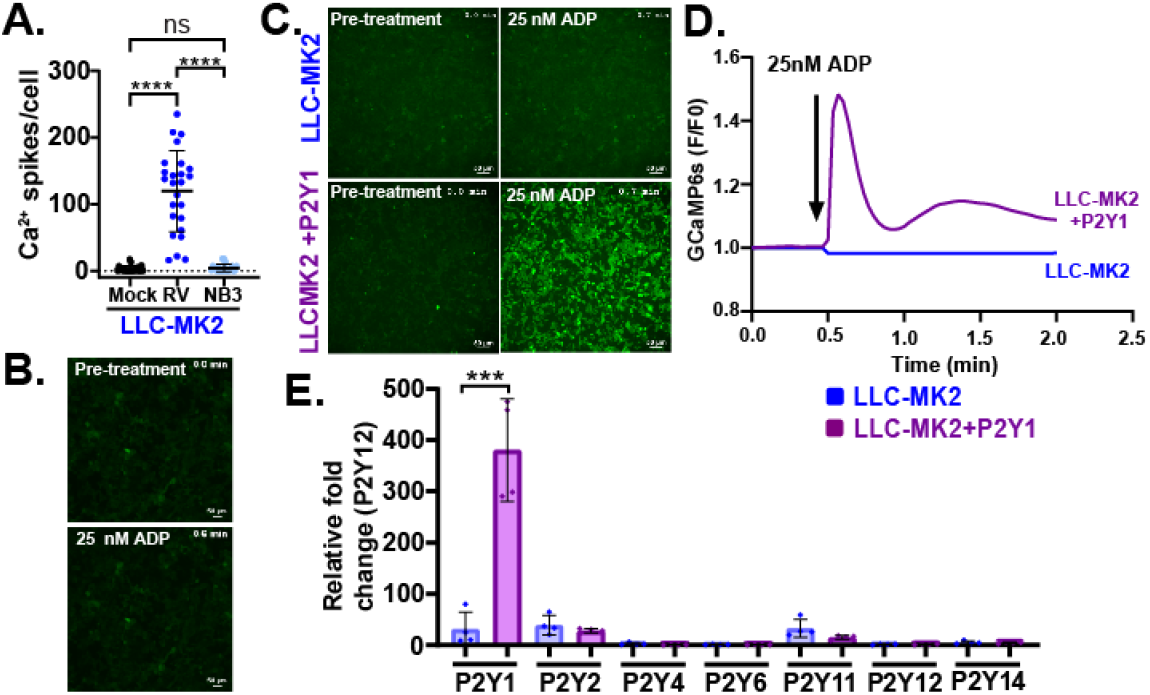
P2Y1 Ca^2+^ signaling in LLC-MK2 cells. **A.** Ca^2+^ spikes per cell in mock (black), rotavirus-infected (blue), and neighboring (NB3) uninfected (light blue) cells. ****p<0.0001 by Kruskal-Wallis with Dunn’s multiple corrections. **B.** Representative images of LLC-MK2 cell monolayers before (top) and after (bottom) treatment with 25 nM ADP. **C.** Representative images of LLC-MK2 and LLC-MK2+P2Y1 cell monolayers before (left) or after (right) agonist treatment with 25nM ADP. **D.** Ca^2+^ signaling traces of LLC-MK2 (blue line) and LLC-MK2+P2Y1 cells (purple line) after treatment with 25 nM ADP (arrow). **E.** P2Y purinergic receptor mRNA expression by qPCR in LLC-MK2 and LLC-MK2+P2Y1 cells. Data were normalized to 18s mRNA and plotted relative to P2Y12 mRNA transcript levels. ***p<0.0001 by individual T tests. All data represented as mean ± SD. Scale bars are all 50 µm.

We next investigated whether exogenous expression of P2Y1 was able to rescue the rotavirus-induced ICWs and, if this affected rotavirus spread. We performed live imaging of rotavirus-infected LLC-MK2-GCaMP6s+P2Y1 cells and found these cells exhibited both higher basal Ca^2+^ signaling and ICWs that originate from rotavirus-infected cells (Figure 8A, Movie S7), resulting in greater Ca^2+^ signaling in the neighboring, uninfected cells than mock-inoculated cells (Figure 8B). Next, we used plaque assays to assess rotavirus spread in LLC-MK2-GCaMP6s cells. In parental LLC-MK2-GCaMP6s cells, rotavirus formed few plaques and those visible were turbid (Figure 8C, top); however, LLC-MK2-GCaMP6s+P2Y1 cells showed more plaques with greater clearing (Figure 8C, bottom). We used fluorescent microscopy to examine spread of SA11-mRuby in the area around plaques for both cell lines. We saw strong mRuby expression for both cell types, but LLC-MK2-GCaMP6s+P2Y1 had significantly more infected cells surrounding the plaques regions with cell monolayer clearing (Figure 8D-E). Thus, introducing P2Y1 expression in LLC-MK2-GCaMP6s cells provides a gain-of-function to produce ICWs in response to rotavirus-infection and this results in better plaque formation and more efficient virus spread. Together these data support our model that the Ca^2+^ signals from P2Y1-mediated ICWs, which requires ER Ca^2+^ release by IP_3_R, and primes neighboring cells to promote more robust rotavirus replication and spread *in vitro*.

**Figure 8.**
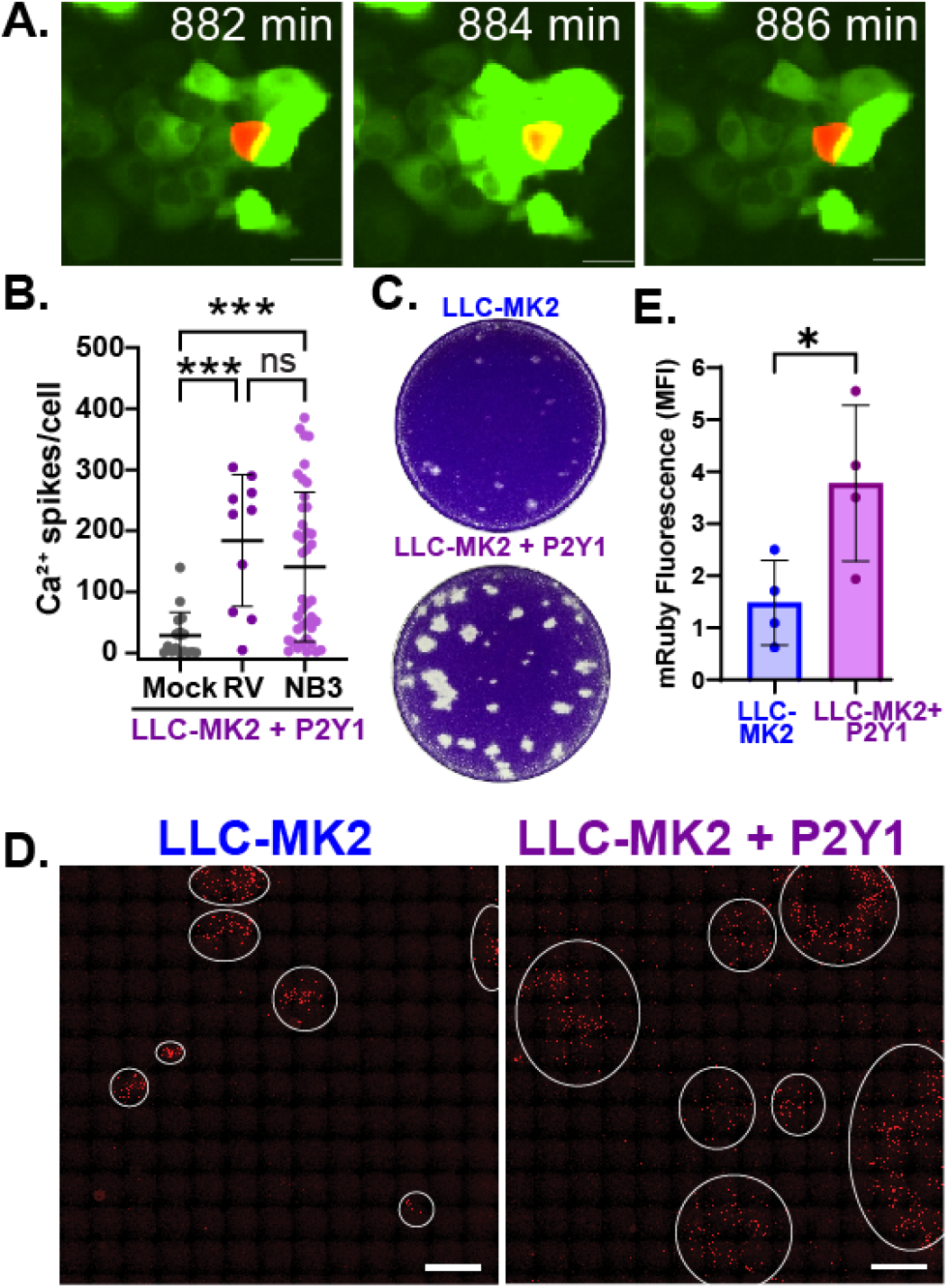
Reconstituting ICWs in LLC-MK2+P2Y1 cells increases rotavirus spread. **A.** Filmstrip images of RV ICWs in LLC-MK2+P2Y1 cells. Scale bar is 50 µm. **B.** Ca^2+^ spikes per cell in mock or RV infected and neighboring (NB3) LLC-MK2 cells. ***p<0.001 by Kruskal-Wallis with Dunn’s multiple correction test. **C.** Representative images of RV plaques formed on LLC-MK2 (Top) and LLC-MK2+P2Y1 (bottom) monolayers. **D.** RV infected (red) LLC-MK2 (left) and LLC-MK2+P2Y1 (right) cell monolayers. White circles designate the margins of RV foci. Images were captured using a Plan Fluor 10x objective (NA 0.30) with 15 x15 mm stich parameters. In these stitched images, the scale bar represents 2000 µm. **E.** Quantitation of mean fluorescence intensity of TRITC channel. *p<0.05 by Mann-Whitney T-test. All data are shown as the mean ± SD.

## Discussion

A hallmark of rotavirus infection, and several other viruses, is an elevation in cytosolic Ca^2+^ and decrease in ER Ca^2+^ stores, which facilitates virus replication and contributes to pathogenesis through a variety of downstream pathways. Host cells also rely on Ca^2+^ signaling pathways to maintain homeostasis. The IP_3_R Ca^2+^ channel is an important Ca^2+^ signaling relay hub converting extracellular signals (e.g., ADP) into intracellular signals in which Ca^2+^ itself is the second messenger that controls a myriad of cellular pathways via Ca^2+^-regulated proteins (30). While the NSP4 viroporin initiates the dysregulation of Ca^2+^ homeostasis during rotavirus infection, the role of IP_3_R in generating rotavirus-induced Ca^2+^ signals, particularly in virus-infected cells, had not been characterized. By developing a MA104-GCaMP6s-IP_3_R-TKO cell line, we uncovered the following: (i) IP_3_R is not required for elevated Ca^2+^ signaling observed in rotavirus-infected cells; (ii) IP_3_R ER Ca^2+^ release is critical for P2Y1-mediated ICWs that increase Ca^2+^ signaling in neighboring, uninfected cells; and (iii) while IP_3_R Ca^2+^ signaling is not *necessary* for rotavirus replication, the rotavirus-induced ICWs, which require IP_3_R, increase the kinetics of rotavirus replication and spread *in vitro*. Together, these findings indicate that increased Ca^2+^ signals in neighboring cells, caused by ICWs, primes these cells to better support rotavirus replication. This process of ‘pre-emptively priming’ uninfected cells within a viral niche represents a novel mechanistic paradigm by which viruses exploit intercellular host responses to promote their replication.

We recently showed that rotavirus-induced increases in cytosolic Ca^2+^ occur through a massive increase in discrete Ca^2+^ signaling events, and a substantial part of this signaling comes from the release of ER Ca^2+^ (20). By examining IP_3_R-null cells (both HEK293 and MA104), we have been able to differentiate two distinct types of Ca^2+^ signals that are induced during rotavirus infection: (1) the intracellular Ca^2+^ signals that occur within rotavirus-infected cells, which were IP_3_R-independent, and (2) the multicellular ICWs that propagate from infected to neighboring, uninfected cells, which are IP_3_R-dependent. By dissecting out these distinct Ca^2+^ signals, we can gain new mechanistic insights into how these signals support rotavirus replication and spread.

The intracellular Ca^2+^ signals are IP_3_R-independent, as knockout of IP_3_R did not reduce the number of Ca^2+^ signals observed. Since NSP4 alone is sufficient to increase Ca^2+^ signaling by release of ER Ca^2+^, we propose that the Ca^2+^ signals observed in the IP_3_R-TKO cells are ER Ca^2+^ release events by the NSP4 viroporin (27, 31); however, some of these Ca^2+^ signals could be Ca^2+^ entry via SOCE channels (e.g., Orai) activated by NSP4-mediated decrease in ER Ca^2+^ levels (28). The elevation in Ca^2+^ signaling, in the absence of IP_3_R, was sufficient to support rotavirus replication, though the kinetics of multi-step virus replication was impaired (discussed below). The fact that rotavirus encodes an intrinsic capability to drive this degree of Ca^2+^ dysregulation further highlights the critical role Ca^2+^ signaling by NSP4, and perhaps other viroporins, plays in virus replication (3, 7, 12). This raises the question of what cellular pathways are activated by these intracellular Ca^2+^ signals. NSP4 viroporin Ca^2+^ signals are critical for rotavirus replication by initiating the cellular autophagy pathway via activation of the Ca^2+^- dependent kinase CaMKKβ (12). Further, these dynamic ER Ca^2+^ signals increase mitochondrial metabolism and suppress the activation of apoptosis, which would also benefit rotavirus replication (20). Yet, unraveling the complexity of virus-induced Ca^2+^ signaling remains a burgeoning field, so we are certainly to discover other Ca^2+^-regulated pathways involved in virus replication.

While IP_3_R Ca^2+^ signaling was dispensable for rotavirus replication, MA104-GCaMP6s-IP_3_R-TKO cells revealed rotavirus induced ICWs significantly increased rotavirus spread and the rate of virus replication. The importance of this paracrine ADP/P2Y1-generated ICW signaling pathway was further supported by rescue of rotavirus plaque formation and increased spread in LLC-MK2-GCaMP6s cells when rotavirus ICW generation was reconstituted by exogenous P2Y1. Further, because MA104-GCaMP6s-IP_3_R-TKO cells still released ADP, we can infer that the loss of IP_3_R Ca^2+^ signaling, and not other downstream effects of P2Y1 activation, cause the defect in rotavirus spread. Since MA104-GCaMP6s-IP_3_R-TKO cells supported replication similar to parental MA104-GCaMP6s cells, these data indicate that IP_3_R increased rotavirus replication and spread via an increase in Ca^2+^ signaling in neighboring, uninfected cells through the P2Y1- mediated ICWs.

Based on these data, we propose the rotavirus-induced ICWs activate pro-viral Ca^2+^- regulated pathways in neighboring cells, ultimately priming these cells for more rapid and/or increased replication. This raises the question of which Ca^2+^-regulated pathways are activated by ICWs in neighboring cells and how do they increase rotavirus replication. One likely pathway is the activation of autophagy because ICWs, like NSP4-mediated Ca^2+^ signals, are generated by release of ER Ca^2+^ and IP_3_R Ca^2+^ release can activate CaMKKβ and AMPK phosphorylation and upregulate autophagy (32-34). In rotavirus-infected cells, early autophagosome membranes are usurped to traffic NSP4 and VP7 out of the ER and form a membrane compartment associated with viroplasms (i.e., virus replication complexes) which is the site of final rotavirus assembly (12). Thus, ICWs could activate the early, biosynthetic stages of the autophagy pathway, making more of these membranes available to be utilized upon rotavirus infection.

Rotavirus-induced Ca^2+^ signaling is also associated with a broad dysregulation in the actin cytoskeleton, though this has primarily been studied in infected or NSP4 expressing cells and the potential role for ICWs has not been investigated (35). Nevertheless, ICWs are an important mode of multicellular epithelial signaling that directs cell extrusion from monolayers in response to cell damage (36-38). The resulting IP_3_R-mediated ER Ca^2+^ release triggers a global, but transient, actin reorganization process termed Ca^2+^-mediated Actin Reset (CaAR) (39). The CaAR response can drive significant changes in host gene transcription by releasing transcription factors otherwise sequestered in the cytosol (39), and this may help identify cellular pathways activated by ICWs. Importantly, the CaAR responses studied thus far have come from singular signals/damage events, but during rotavirus infection, neighboring cells are stimulated by hours of ICWs, providing ample time and numbers of signals to drive substantial changes. Thus, it will likely require a detailed, multi-omics approach to identify ICW-response pathways and elucidate which of those support rotavirus spread.

In summary, we have uncovered a dichotomous role for IP_3_R in the overall Ca^2+^ signaling landscape during rotavirus infection and in rotavirus replication and spread. Within infected cells, rotavirus (presumedly via NSP4) generates sufficient Ca^2+^ signaling to support its replication without IP_3_R, making this host channel dispensable. In contrast, rotavirus spread was increased by the presence of ICWs and therefore increased Ca^2+^ signaling in neighboring cells, essentially priming them for future rotavirus infection. This implies that viral take-over of the infection niche goes beyond rotavirus-infected cells to include nearby uninfected cells, which undergo “preemptive reprogramming” by repeated ICWs prior to becoming infected. Further, ICWs triggered by rotavirus infection may be a common host response to many different virus infections, and, if so, this P2Y1-mediated Ca^2+^ signaling pathway would have a broader importance in virus replication and pathogenesis. Finally, purinergic signaling is just one of the myriad of other intercellular signaling molecules/pathways that could be exploited by viruses to prime or otherwise reprogram uninfected cells within the infection niche. Identification of analogous virus-induced intercellular signaling pathways may uncover new mechanisms by which viruses, or other microbes, exploit host responses to benefit their replication and spread.

## Acknowledgments

We thank Dr. David I. Yule at University of Rochester Medical School for his expertise and advise in generating and characterizing the MA104-GCaMP6s-IP_3_R-TKO cells. This work was supported in part by NIH grants R01DK115507 and R01AI158683 (PI: J. M. Hyser). Trainee support for J.L.P. was provided by NIH grants F31AI169983 (PI: J. L. Perry). Trainee support for F.J.S. was provided by NIH grants F31DK132942 (PI: F. J. Scribano). Trainee support for J.T.G was provided by NIH grants F30DK131828 and Histochemical Society Graduate Medical Trainee and Graduate Student Cornerstone Grant (PI: J.T. Gebert). Trainee support for K.A.E. was provided by NIH grants F32DK130288 and Histochemical Society Postdoctoral Keystone Grant. Lentivirus packaging of the P2Y1 expression construct was supported by the ATC Gene Vector Core at Baylor College of Medicine.

